# Detecting anomalies in RNA-seq quantification

**DOI:** 10.1101/541714

**Authors:** Cong Ma, Carl Kingsford

## Abstract

Algorithms to infer isoform expression abundance from RNA-seq have been greatly improved in accuracy during the past ten years. However, due to incomplete reference transcriptomes, mapping errors, incomplete sequencing bias models, or mistakes made by the algorithm, the quantification model sometimes could not explain all aspects of the input read data, and misquantification can occur. Here, we develop a computational method to detect instances where a quantification model could not thoroughly explain the input. Specifically, our approach identifies transcripts where the read coverage has significant deviations from the expectation. We call these transcripts “expression anomalies”, and they represent instances where the quantification estimates may be in doubt. We further develop a method to attribute the cause of anomalies to either the incompleteness of the reference transcriptome or the algorithmic mistakes, and we show that our method precisely detects misquantifications with both causes. By correcting the misquantifications that are labeled as algorithmic mistakes, the number of false predictions of differentially expressed transcripts can be reduced. Applying anomaly detection to 30 GEUVADIS and 16 Human Body Map samples, we detect 103 genes with potential unannotated isoforms. These genes tend to be longer than average, and contain a very long exon near 3′ end that the unannotated isoform excludes. Anomaly detection is a new approach for investigating the expression quantification problem that may find wider use in other areas of genomics.

## 1 Introduction

While modern RNA-seq quantification algorithms [e.g. 1–7] often achieve high accuracy, there remain situations where they give erroneous quantifications. For example, most quantifiers rely on a predetermined set of possible transcripts; missing or incorrect transcripts may cause incorrect quantifications. Read mapping mistakes and unexpected sequencing artifacts introducing technical biases also lead to misquantifications. Incomplete sequencing bias models can mislead the probability calculation of which transcripts generate the reads. Quantification algorithms themselves could introduce errors since their objectives cannot typically be guaranteed to be solved optimally in a practical amount of time.

When interpreting an expression experiment, particularly when a few specific genes are of interest, the possibility of misquantification must be taken into account before inferences are made from quantification estimations or differential gene expression predictions derived from those quantifications. Expression quantification is the basis for various analyses, such as differential gene expression [8], co-expression inference [9], disease diagnosis and various computational prediction tasks [e.g., 10–12]. Statistical techniques such as bootstrapping [13] and Gibbs sampling [1, 14, 15] can associate confidence intervals to expression estimates, but these techniques provide little insight into the causes of low confidence or misquantification, and detect a subset of misquantifications.

Here, we introduce a method to identify potential misquantifications using a novel anomaly detection approach. This approach automatically identifies regions of known transcripts where the observed fragment coverage pattern significantly disagrees with what the coverage is expected to be. These regions indicate that something has gone “wrong” with the quantification for the transcripts containing the anomaly: perhaps a missing transcript, missing features in the probabilistic model, an algorithmic failure to optimize the likelihood, or some other unknown problem.

One advantage of this model-based anomaly detection approach is that it does not require any known ground truth to discover potential errors. The expected and observed coverages are intermediate values in the quantifier. The expected coverage is derived from a bias correction model that is used by modern RNA-seq quantification algorithms to model fragment generation biases with varied GC content, sequence, and position in the transcript [7, 16]. In order to take into account other aspects of sequencing (such as read mapping quality, fragment length distribution), quantifiers sometimes cannot assign fragment in proportion to the expected coverage. By comparing the expected and observed coverages, anomaly detection identifies cases where it is not possible to satisfy the assumed model of fragment generation.

Another advantage of our proposed anomaly detection method is that it can provide more insight into what is causing the misquantification by identifying specific regions of specific transcripts for which the assumed theoretical model of read coverage does not match what is observed. These anomaly patterns can then be used to derive hypotheses about the underlying cause. For example, systematic lower-than-expected expression across an exon may indicate the existence of a unknown isoform that omits that exon. In this way, anomalies are more informative and suggestive of the cause of misquantifications than confidence intervals.

A third advantage of the approach is that the anomalies can be used to design better quantification algorithms. When there is good reason to believe the transcriptome annotations and sequencing is of high quality, analyzing the cause of anomalies could reveal new features or aspects of the sequencing experiment that may improve the quantification model, and may therefore be used to inspire improvements to, e.g., bias correction models or optimization approaches.

Anomaly detection has been applied to other areas in genomics where it has proved its usefulness. In genome assembly, anomaly detection has been used to detect low-confidence assembled sequences. Genome assembly algorithms seek a set of sequences that can concordantly generate the WGS reads and can be assumed to have near uniform coverage. The assembled sequences that do not fit this assumption can be hypothesized to contain errors and have low reliability [17]. Similarly, anomaly detection in transcriptome assembly identifies unreliable transcript sequences [18]. Low-confidence assembly detection has been used to analyze non-model organisms and incorporated into analysis workflows [19–21].

In RNA-seq expression quantification, some research has been conducted on how to identify anomalous predictions. For example, Robert and Watson [22] identify uncertainties in gene-level quantification related to gene sequence similarity by comparing against an external ground truth. However, uncertainties do not necessarily indicate anomalous quantification. In addition, external information about sequence similarity provides limited insight on how to improve the quantification models. Soneson et al. [23] use a compatibility score of observed and predicted junction coverage to indicate genes with potential misquantification in its transcripts. With this anomaly score, it is possible to narrow down the misquantified transcripts by the anomalous splicing junctions. Nevertheless, it does not directly spot the misquantified ones, nor does it predict the potential cause of the low quantification reliability.

In this work, we detect quantification anomalies using the disagreement between the modeled expected distribution and the observed fragment coverage distribution that is obtained after the quantifier has allocated fragments to transcripts. We do this by introducing an anomaly metric to quantify regions of high disagreement. Specifically, we identify the contiguous regions that have the largest difference between these two distributions. This metric has the natural biological meaning as the largest over-or under-expression (compared with what is expected) of any region within the transcript. We further begin to categorize the anomalies by their causes: adjustable anomalies are the ones possibly caused by quantification algorithm mistakes, and unadjustable anomalies are those possibly caused by transcripts missing from the reference transcriptome. This categorization is done by correcting quantification deviations using a fragment reassignment procedure based on linear programming (LP) to attempt to correct anomalies. Those anomalies that can be corrected this way are candidates for having been caused by algorithmic error. The fragment reassignment procedure also generates an adjusted abundance estimation to when correcting the quantification of the adjustable anomalies.

Because it includes a rich bias model, we use Salmon [7] as the base quantifier on which to build and test anomaly detection, and we term our implementation Salmon Anomaly Detection (SAD). However, the idea of anomaly detection can be applied to any method that generates an internal model of expected sequence coverage.

Applied to 30 GEUVADIS [24] samples and 16 Human Body Map [25] samples, SAD identifies both adjustable and unadjustable anomalies. The anomalous transcripts often have a different set of protein domains from other isoforms in the same gene or belong to cell type marker genes. For example, a kidney cell type marker gene *TAX1BP3* has an adjustable anomalous in one of its transcripts, suggesting a change read assignment across isoforms should be made. An isoform of the gene *UBE2Q1* is identified to be an unadjustable anomaly, and the isoform is the only one in the gene to contain the ubiquitin-conjugating enzyme domain. Using the adjusted abundance estimates corresponding to the adjustable anomalies, the number of falsely detected differentially expressed transcripts can be reduced by 2.4% – 6% in the GEUVADIS samples.

We observe some common patterns of the unadjustable anomalies that are shared among all GEUVADIS and Human Body Map samples: genes containing the anomalies tend to be longer than average, and contain a long exon at the 3′ end. The hypothesized unannotated sequences tend to have an early transcription stop in the middle of the 3′ long exon. But we are not sure about the cause of generating the unannotated sequences, nor their functions.

We further validate SAD’s prediction via simulation and show that, both adjustable and unadjustable anomalies of SAD precisely describe the corresponding types of misquantification. The read re-assignment procedure of SAD generates an adjusted quantification that is closer to the simulated expression and reduces the mean ARD distance by about 0.05. Surprisingly, in simulation, unadjustable anomalies reflect the existence of novel isoforms with 3% – 35% higher precision compared with applying transcriptome assembly to the samples, when the novel isoforms contain alternative starting / ending sites.

## 2 Results

### 2.1 Overview of anomaly detection and categorization

SAD defines transcripts with anomalous read coverage (Figure 1) as those for which the observed coverage distribution contains a significantly over-expressed or under-expressed region compared to the expected coverage (Section 4.1). Both the observed and the expected distribution are calculated by the Salmon quantifier [7]. The observed distribution is the weighted number of reads assigned to each position in the transcript as processed by Salmon (Section 4.5). The expected distribution estimated by Salmon is the probability of generating a read at each position considering the surrounding GC content, K-mers, and the position in the transcript (Section 4.4). The anomaly metric can be confounded by either a low expression abundance or an estimation error of the expected distribution. To remove the confounding effect, we model the anomaly metric probabilistically (Section 4.2) and use the empirical p-value to determine whether the observed difference is statistically significant and whether the transcript should be labeled as an anomaly (Section 4.3).

**Figure 1:**
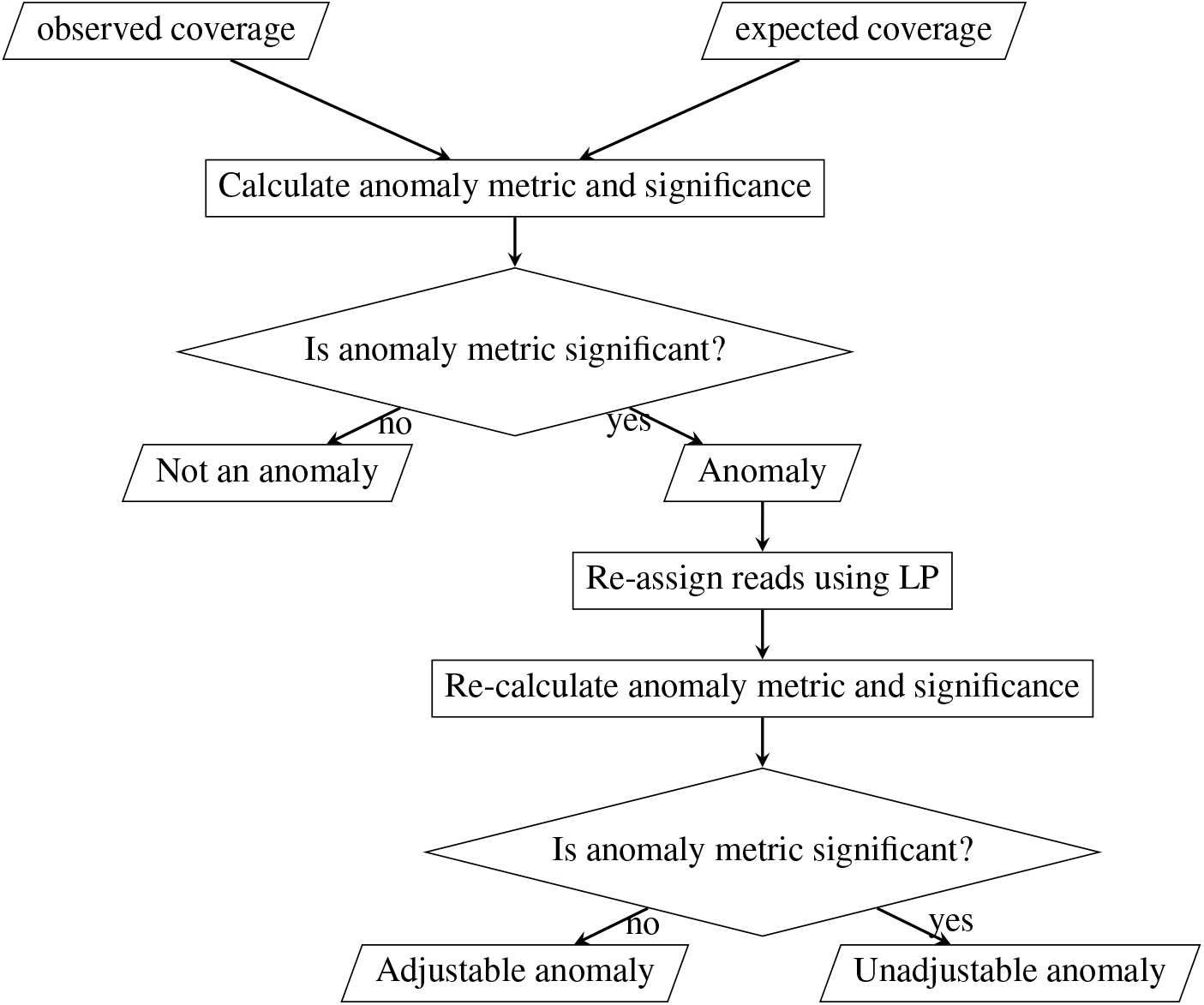
Diagram of SAD. SAD detects anomalies by calculating an anomaly metric and the significance of its value. To further distinguish the potential cause of the anomalies, it re-assigns the reads across isoforms and checks whether the anomaly metric becomes insignificant after re-assignment. The anomalies whose anomaly metrics become insignificant are categorized as adjustable anomalies, and considered to be caused by quantification algorithm mistake. The anomalies whose anomaly metrics remain significant are categorized as unadjustable anomalies, and considered to be caused by the unannotated transcript sequences, that is, the incompleteness of the reference transcriptome.

Anomaly categorization is done by re-assigning the reads across the isoforms using linear programming (LP) (Section 4.6) and checking whether the anomaly metric becomes insignificant after the re-assignment. Though it is possible that reads are mapped to multiple genes, the majority of reads are multi-mapped to isoforms within the same gene, and the re-assignment is performed within each gene to control the size of the LP. If the anomaly metric remains significant after the re-assignment, the anomaly is labeled as an unadjustable anomaly and considered to be caused by the incompleteness of the reference transcriptome. Otherwise, it is labeled as an adjustable anomaly and is potentially caused by quantification algorithm error.

SAD gives rise to two outputs: (1) a list of unadjustable anomalies and (2) the transcript-level adjusted quantification for the genes containing the adjustable anomalies. Each output has a direct application: the unadjustable anomalies can be used as a predictor for novel isoforms; the adjusted quantification can replace Salmon’s quantification for the adjusted subset of transcripts and be used in any analyses depending on RNA-seq quantification.

### 2.2 Examples of detected anomalies

We provide some examples of the detected anomalies after applying SAD to 30 GEUVADIS [24] and 16 Human Body Map datasets [25]. The 30 GEUVADIS samples are the same as in the work of Patro et al. [7], in which 15 lymphoblastoid cell lines from the Toscani in Italia (TSI) population are sequenced, where each cell line is sequenced twice, at two different sequencing centers. The Human Body Map project data consists of 16 samples each from a different tissue, including adrenal, adipose, brain, breast, colon, heart, kidney, liver, lung, lymph, ovary, prostate, skeletal muscle, testes, thyroid, and white blood cells. The Pfam annotation [26] is used to label protein domains. Among the examples of anomalies, some contain protein domains that are different from other isoforms, and some belong to cell type marker genes.

SAD identifies an adjustable anomaly in the gene *TAX1BP3* in the kidney sample from the Human Body Map dataset. The *TAX1BP3* gene is potentially a cell type marker gene for podocytes cells in kidney [27]. One isoform (ENST00000611779.4) of this gene has an under-expression anomaly in the first 200 bp (Figure 2A). This under-expression anomaly can be adjusted by re-assigning reads between this and another isoform, ENST00000225525.3 (Figure 2B). The expression estimations are changed according to the adjustment: the abundance ratio between these two isoforms decreases from 4.5 to 0.9. The difference between the two isoforms is that the second exon of ENST00000225525.3 is excluded in ENST00000611779.4 (Figure 2C). This exon is located in the middle of the PDZ domain, the function of which is to help protein scaffolding and receptor anchoring. Having a more accurate quantification of the two isoforms can be important in analyzing the effect of the middle exon using expression.

**Figure 2:**
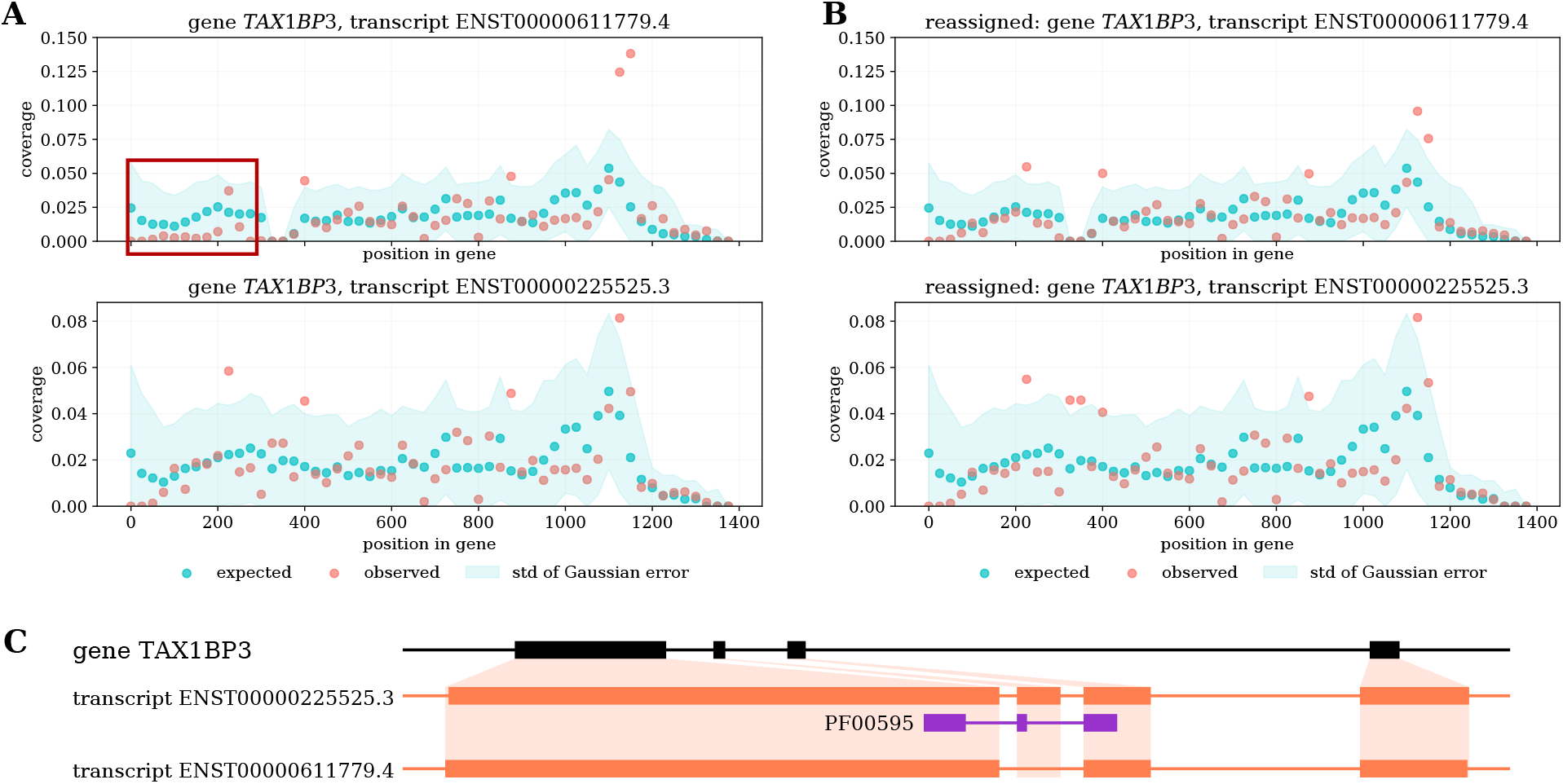
Adjustable anomalies in the kidney sample of the Human Body Map dataset. (A)Red and blue points are the observed and expected coverage distribution before SAD adjustment. The anomaly transcript ENST00000611779.4 has an under-expression in the first 200 bp (top), marked by the red box. Another transcript is involved in the adjustment (bottom). (B) The distributions of the same pair of transcripts after SAD adjustment. (C) The protein domain annotation of the two transcripts.

Another example of an adjustable anomaly is within *BIRC3* gene in one GEUVADIS sample. This gene is involved in apoptosis inhibition under certain conditions. The second half of the isoform ENST00000532808.5 is under-expressed under the read assignment of Salmon (Figure 3A). Re-assigning the reads between this isoform and another isoform ENST00000263464.7 removes the under-expression phenomenon (Figure 3B), and at the same time alters the expression level of both isoforms. The original expression abundances were similar to each other, but after SAD adjustment ENST00000263464.7 has 3 times the expression of ENST00000532808.5. The two isoforms are different in their starting and ending positions but have the same set of internal exons. The protein domains between the two isoforms are the same according to Pfam annotations (Figure 3C). Nevertheless, a better read assignment can improve the normalized abundance estimation of the whole gene.

**Figure 3:**
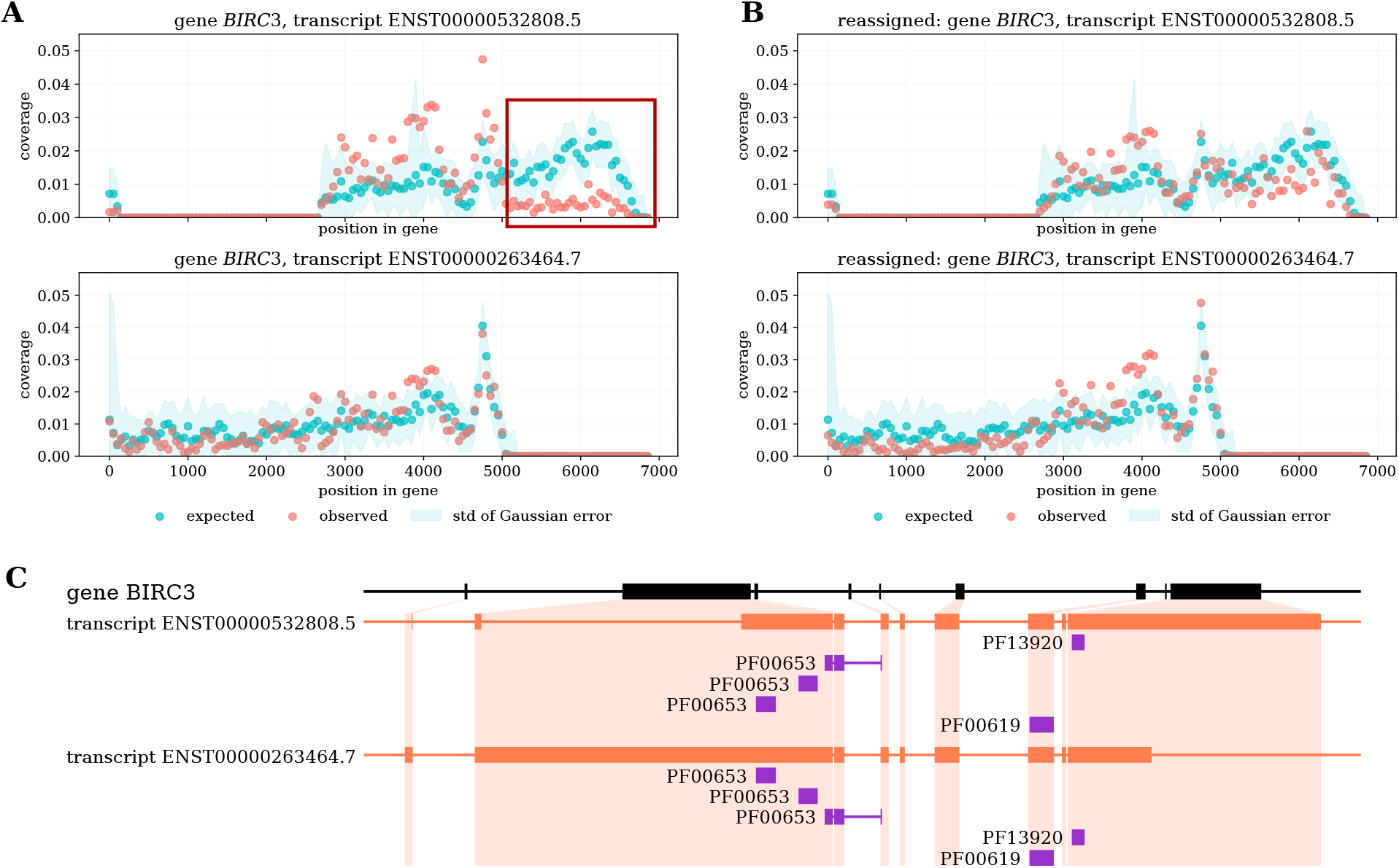
Adjustable anomalies in a sample of GEUVADIS dataset. (A) The top transcript ENST00000532808.5 is identified to be adjustable anomaly, and its under-expression anomaly region is marked by the red box. The bottom transcript is involved in the quantification adjustment. (B) The observed and expected distribution after SAD adjustment. (C) The protein domain annotation of the previous transcripts.

SAD also reveals unadjustable anomalies in isoforms that have a different set of protein domains from the other isoforms of the same gene. For example, gene *UBE2Q1* gene and gene *LIMD1* in the heart sample of the Human Body Map dataset contain unadjustable anomalies (Figure 4), suggesting the existence of unannotated isoforms. In both genes, the protein domains in the anomalous isoform are different from those in the other annotated isoforms: ENST00000292211.4 of gene *UBE2Q1* is the only annotated isoform that has ubiquitin-conjugating enzyme domain, and ENST00000273317.4 of gene *LIMD1* contains three zinc-finger domains while the other isoforms only contain two or zero. The unannotated novel sequences of both genes potentially have the same set of protein domains as the anomalies, as suggested by the over-expression region. The Scallop transcript assembler [28] is able to assemble a novel sequence of *LIMD1* without the under-expression region, thus supporting this detected anomaly.

**Figure 4:**
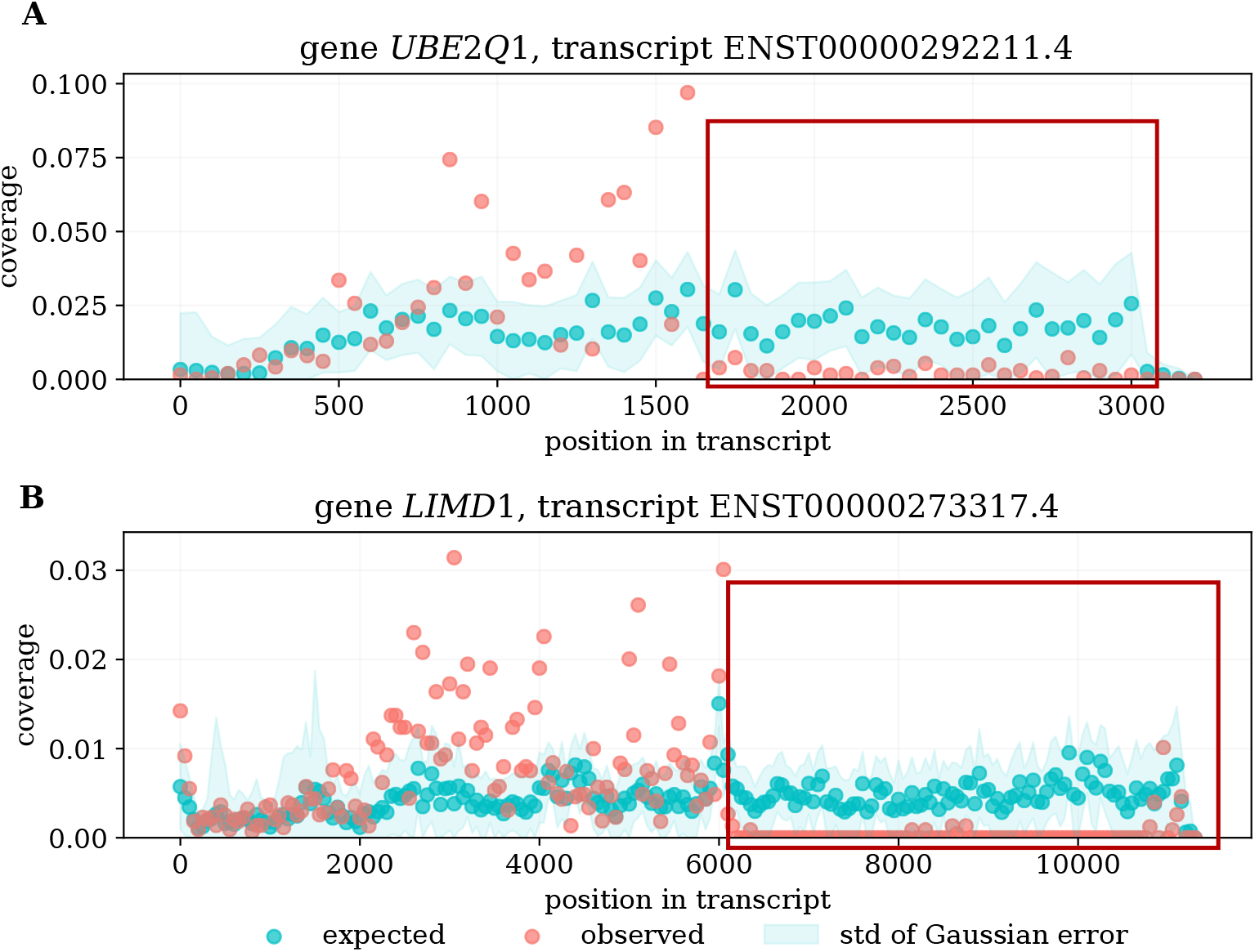
Examples of unadjustable anomalies identified by SAD. (A) An anomaly isoform of gene *UBE2Q1* (B) An anomaly isoform of gene *LIMD1*. Both examples are found in the heart sample of the Human Body Map dataset. Red and blue points are the observed and expected coverage distribution of the anomaly transcripts, and the blue shade is the standard deviation of the expected distribution estimation. The red box indicates the under-expression anomaly region. For both genes, the transcript region near 5′ end is over-expressed, and the region near 3′ end is under-expressed.

### 2.3 Adjustable anomalies give an adjusted quantification that reduces false positive differential expression detections

The adjusted quantification of SAD reduces the number of false positive calls in detecting differentially expressed transcripts. Previously, Patro et al. [7] showed that the 30 TSI samples from GEUVADIS dataset [24] likely do not have differential expressed transcripts, but quantification mistakes can lead to false positive differential expression (DE) predictions across sequencing center batches. They also showed that a more accurate quantification can reduce the number of false positive detections. We apply SAD to the same samples and compare the number of differentially expressed transcripts detected using Salmon’s original quantification and SAD-adjusted quantification. SAD-adjusted quantification uses SAD-adjusted estimates for the adjustable anomalies and other isoforms involved in read re-assignment, but uses the Salmon [7] quantification elsewhere. Differential expression is inferred by DESeq2 [29] on the transcript level. On this data, the number of DE transcripts is reduced by about 2.4% – 6% with various FDR threshold when using SAD-adjusted quantification compared to Salmon’s original quantification (Table 1). This provides evidence that these anomalies are likely true errors, that are correctable using a different read re-assignment procedure from Salmon.

**Table 1:** Number of DE transcripts detected at given FDR threshold. Among the 30 samples, there should not be any DE transcripts. With SAD-adjusted expression quantification, the number of false positively detected DE transcripts is reduced.

| FDR | Salmon | SAD-adjusted | percentage reduced |
| --- | --- | --- | --- |
| 0.01 | 6088 | 5723 | 6.00% |
| 0.05 | 10132 | 9777 | 3.50% |
| 0.1 | 13555 | 13228 | 2.41% |

### 2.4 Genes that contain common unadjustable anomalies tend to be long and have long exons

There are unadjustable anomalies common to all GEUVADIS and Human Body Map dataset, and the genes containing them tend to have a larger gene length than average (Figure 5A). There are 103 common genes containing unadjustable anomalies in all 30 samples in GEUVADIS and 16 samples in Human Body Map (see Supplementary Table for the full list). These genes span 23 chromosomes and contain various number of annotated isoforms ranging from 1 to 24. One common pattern about these genes is that they tend to be long, and the existence of unadjustable anomalies may be related to the gene structure.

**Figure 5:**
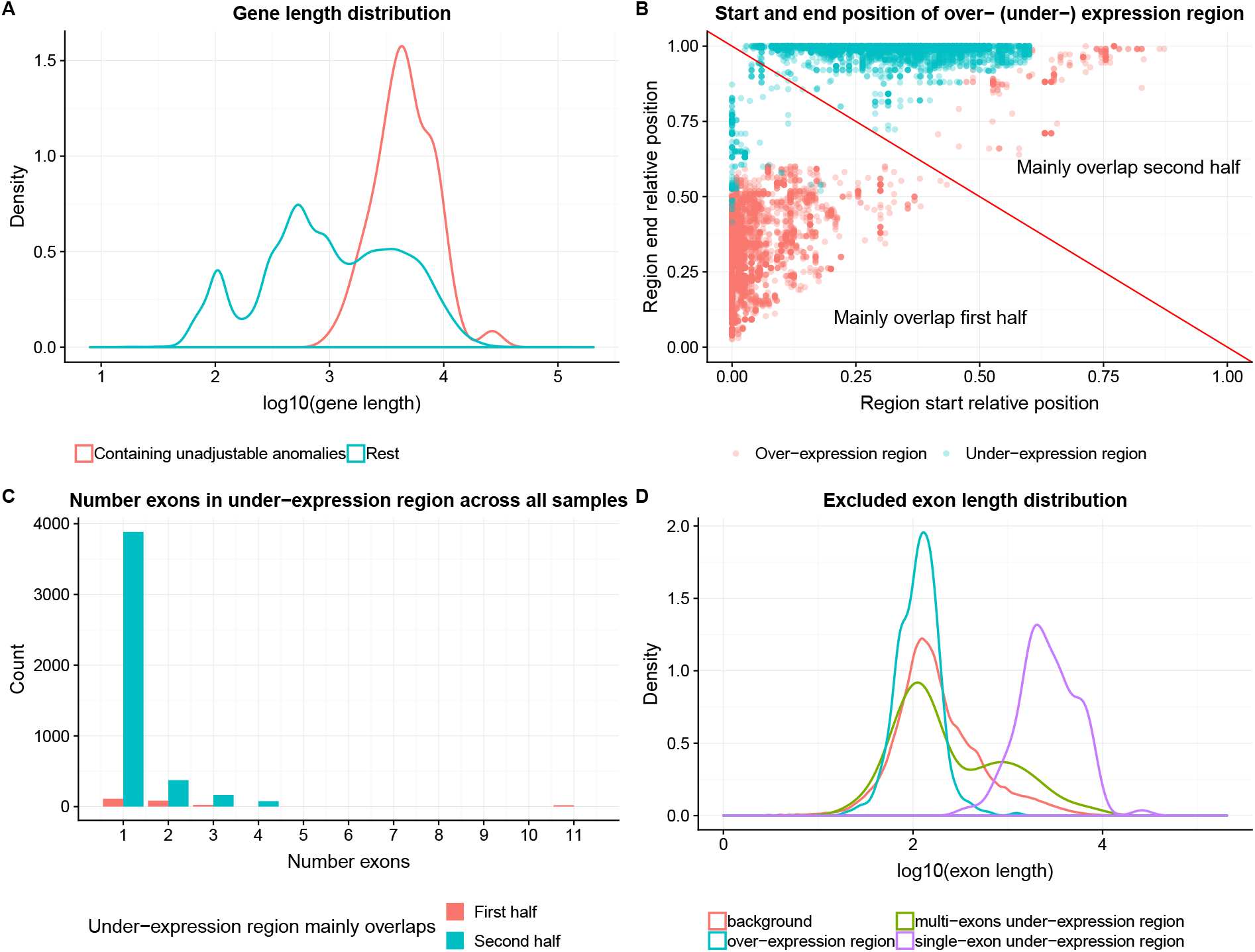
(A) Density curve of gene length distribution of unadjustable-anomaly-containing genes and the rest. Genes containing unadjustable anomalies tend to be long genes. (B) The start and end proportion of the over-expression and under-expression region of anomalous transcripts of the common unadjustable-anomaly-containing genes. The red diagonal line separates between anomalies of which the over- (under-) expression regions mainly overlap with the first half (5′ half), and the second half (3′ half) of the transcripts. For most of the anomalies, the over-expression region mainly overlaps with the first half of the anomalous transcript, and the under-expression region mainly overlap with the second half of the anomalous transcript. (C) Histogram of number of exons spanning under-expression region of the anomalies corresponding to the 103 shared genes. The count of y-axis is summed over all 46 samples. The under-expression region usually only contain one or a partial exon. (D) Exon length distribution of the exons contained in the over-expression region, under-expression region, and the background of all annotated exons. Two curves are plotted for the exons spanning the under-expression region, corresponding to the case when under-expression regions are single-exon-spanning, and when it spans more than one exons. For the single-exon-spanning underexpression region, the exon tends to be much longer than average.

For most anomaly transcripts of these common genes, the over-expressed regions tend to mainly overlap with the first half of the transcripts near the 5′ end (Figure 5B). Correspondingly, the under-expressed regions are usually located towards the second half of the transcripts near the 3′ end. The under-expression anomaly regions usually only span one exon or a partial exon (Figure 5C). This tends to be true no matter whether their under-expression region mainly overlaps the first half or second half of the transcript. This suggests that there possibly exist unannotated transcripts that have the same intron chain but different transcript starting and ending locations from the known ones. We use single-exon-spanning region to refer to the region only spanning one or a partial exon.

For the single-exon-spanning, under-expressed regions, the exon length distribution is shifted longer compared with the background exon length distribution (Figure 5D), where the background includes all annotated exons. For comparison, we compute exon length distributions of two other types of exons: the exons contained in the over-expressed regions and the exons in under-expressed regions when the regions cover more than one exon. For both of these two types of exons, the length distributions are similar to the background. Considering that the single-exon-spanning, under-expressed regions are the majority of all under-expressed regions (Figure 5C), we conclude the under-expression anomaly often occurs when there is a very long exon and when the long exon is near 3′ end of the transcript.

About 50%–60% of the detected unadjustable anomalies have a corresponding novel isoform assembled by transcriptome assembly algorithms, specifically StringTie [30] and Scallop [28] (Supplementary Figure S1). (See Supplementary Text for the detail of running transcriptome assembly software.) An assembled isoform corresponds to a predicted unadjustable anomaly if the assembled isoform contains all the splicing junctions within the over-expressed region and excludes at least half of the under-expressed region.

Meanwhile, there are 40%–50% of the unadjustable anomalies that do not have a corresponding isoform assembled by transcriptome assemblers. Assuming the expected coverage distribution is modeled correctly, these unadjustable anomalies are likely to indicate true novel isoforms that are not able to be detected by transcriptome assemblers. This is somewhat surprising, but not entirely unexpected given the low overall sensitivity of transcript assembly methods.

While we hypothesize that the unadjustable anomalies are caused by the existence of unannotated transcripts, it is not clear whether the unannotated ones are natural, well-functioning novel isoforms, or non-functioning sequences due to errors in transcription that terminates transcription early in long exons at the 3′ end.

### 2.5 Simulation supports the accuracy of SAD for detecting and categorizing anomalies

On simulation data, both unadjustable and adjustable anomalies precisely reflect the misquantification due to those corresponding causes. We created 24 datasets by varying the number of simulated novel isoforms, the gene annotations, and the expression matrices. (See Supplementary Text for the details of the simulation procedure.)

The unadjustable anomalies reflect the simulated novel transcript sequences with 3%–35% higher precision compared to transcriptome assembly methods (Figure 6A, Supplementary Figure S2A). (See Supplementary Text for the detail of running transcriptome assembly software.) In this comparison, the precision is calculated only for novel isoforms without new splicing junctions, in which case transcriptome assembly could not use accurate spliced alignment to detect novel isoforms. We consider the following two types of assembled transcripts as novel isoforms: (1) the intron chain of the assembled transcript does not exactly match the intron chain of any existing transcript; (2) the intron chain exactly matches one existing transcript, but either transcript starting position or stopping position is more than 200 bp away from the matched existing transcript. Transcriptome assembly methods tend to reconstruct novel isoforms for far more genes than the anomolies detected by SAD. To compare the precision on the same ground, we select the same number of predictions for transcriptome assembly and SAD by selecting those assembled transcripts with the highest coverage. The main advantage of SAD is precision, but not sensitivity, because not not all unannotated isoforms will significantly alter the coverage of known ones (Supplementary Figure S2B). The higher precision of SAD is possibly due to the accurate expected distribution used by Salmon, whereas transcriptome assembly methods usually assume a uniform coverage in the algorithms. When the simulated novel isoforms do not contain new splicing junctions, the coverage is the main indicator of new starting or ending sites of the isoform. In this case, SAD is able to detect the novel isoforms more precisely than transcriptome assembly methods by taking advantage of the accurate expected distribution.

**Figure 6:**
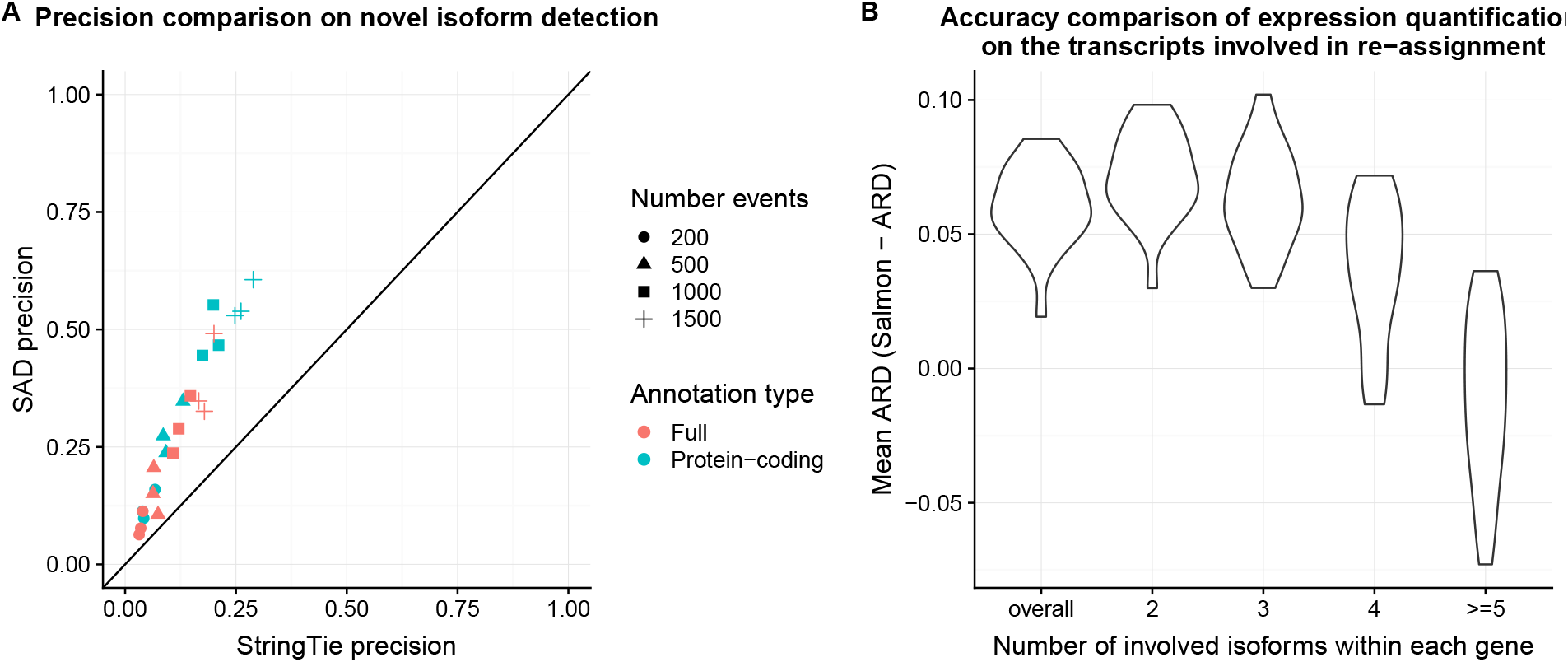
(A) Precision of novel isoform detection of SAD and StringTie. Point color and shape refers to different simulation settings. With the refined expected distribution, the unadjustable anomalies in SAD reflect the simulated novel isoforms more precisely compared to transcriptome assembly method, StringTie. The simulated novel isoforms do not contain new splicing junctions, but only contain new starting / ending sites, or new combinations of known splicing junctions. (B) Quantification accuracy improvement of SAD compared to original Salmon. Each violin refers to a subset of transcripts where the corresponding gene contain a certain number of isoforms in the adjustment according to the x-axis. “Overall” in the x-axis is the overall mean ARD improvement of all adjusted isoforms without distinguishing the number of isoforms involved. The improvement decreases as the number of involved isoforms increases, possibly because the estimation error in the expected distribution is magnified when the LP coefficient grows large.

In addition, the adjusted quantification of SAD is more accurate compared to the original Salmon quantification [7] (Figure 6B, Supplementary Figure S3) on simulation data. SAD is able to reduce the level of misquantification for adjustable anomalies. The accuracy of quantification is measured by the mean ARD (absolute relative difference) [7] between the quantification and the simulated ground truth. ARD is the absolute difference between the estimation and the true value, normalized by the sum of the estimation and the true value. A smaller value of mean ARD indicates an estimator that is closer to the ground truth. However, the accuracy improvement of SAD decreases as more isoforms of one gene are involved in the quantification adjustment. The decrease of improvement is possibly because the estimation error in the expected distribution is magnified when the LP coefficient matrix used by SAD is large in size and potentially ill-conditioned. When the coefficient matrix is ill-conditioned in the linear system, the output can greatly change even with a small error in the input.

## 3 Discussion

We present Salmon Anomaly Detection (SAD), an anomaly detection approach to identify misquantification of expression. SAD detects anomalies by comparing the expected and the observed coverage distribution, and calculating the significance of the over- or under-expression. SAD also categorizes the anomalies into adjustable anomaly and unadjustable anomaly categories to indicate two possible causes of misquantifications: algorithmic errors and reference transcriptome incompleteness. The categorization is done by re-assigning reads across isoforms to minimize the number of significant anomaly scores. We show on simulation data that the detected anomalies and the categorization is reasonable: the unadjustable anomalies predict the existence of novel isoform with higher precision than transcriptome assembly methods, and the read re-assignment leads to adjusted quantification that is closer to the simulated ground truth compared to the original quantification.

Applying SAD on GEUVADIS and Human Body Map datasets, we are able to identify adjustable and unadjustable anomalies that affect isoforms with different protein domains from other isoforms and isoforms from cell type marker genes. Using the adjusted quantification associated with the adjustable anomalies, the number of false positive predictions of differentially expressed transcripts can be reduced. There are common unadjustable-anomaly-containing genes across all samples. Most of the common unadjustable anomalies have an under-expressed region towards the 3′ end of the transcript. The genes that contain the common unadjustable anomalies tend to be longer in length, and their 3′ exons tend to be longer than average.

SAD is only able to detect the subset of misquantifications that have a distorted observed coverage from the expected one. However, some misquantifications may not alter the shape of the observed coverage distribution. For example, high sequence similarity between a pair of transcripts can also lead to severe mis-quantification, however, the read coverage can be close to the expectation for both. Alternatively, the coverage distribution of a lowly expressed existing isoform can be affected by a lowly expressed novel isoform. In this case, the p-value of the anomaly score may not be significant due to the large fluctuation of the observed coverage due to the low expression. Developing other metrics, for example, using transcript similarity or discordant read mapping, could potentially increase the sensitivity and the types of possible misquantification of detection.

For novel isoform detection, only the existence is predicted by SAD, not the sequence or exon-intron structure of the novel isoforms. Retrieving the exon-intron structure remains a problem. Simply combining the existence prediction of SAD with the assembled sequences from transcriptome assembly does not solve the problem of reconstructing novel isoform sequences. About 40%–50% of SAD’s predictions are not assembled by transcriptome assembly methods in the GEUVADIS and the Human Body Map datasets. Incorporating the expected coverage distribution in transcriptome assembly may be a direction to predict the exact exon-intron structure of the novel isoforms.

An improvement in the accuracy of the approximation of the expected distribution may further increase the accuracy in novel isoform prediction and re-quantification by SAD. Currently, the expected distribution is approximated by a bias correction model that uses GC, sequence, and position biases. However, the sequence bias may also be affected by secondary structure of cDNA, which is not considered in current modeling of biases. Additionally, different subtypes of biases can be coupled together, meaning that inferring each type of bias separately may not be sufficient.

SAD takes about eight hours to run on each RNA-seq sample using eight threads. The long running time is mainly due to the sampling procedure in the empirical p-value calculation for all transcripts. A derivation of a p-value approximation to avoid sampling could potentially decrease the resource requirement for computation. Implementation tricks and engineering can also applied to reduce the running time, however this is out of the scope of this work.

Our formulation of anomaly detection is an example of algorithmic introspection: algorithms that can automatically identify where their predictions do not fit the assumptions of the algorithm. This type of algorithmic reasoning is likely to become even more useful as the sophistication of bioinformatics analysis tools increases.

## 4 Methods

### 4.1 An anomaly detection metric

#### Definition 4.1

(Expected coverage distribution). Given transcript *t* with length *l*, and a fragment *f* that is sequenced from *t*, the starting position of *f* is a random variable with the possible positions {1, 2, 3, ⋯, *l*} as its domain. The expected coverage distribution of *t* is the probability distribution of the starting position of any fragment *f*. The expected coverage distribution for each transcript *t* sums to 1.

With a non-zero fragment length, the viable starting position excludes the last several positions in the transcript. The probability of the last several positions in the expected coverage distribution is set to 0 to account for the fact that they are not viable. After aligning and assigning the sequencing reads to transcripts, the number of fragments starting at each position can be observed and counted; this is referred to as the observed coverage. The observed coverage can be converted to a distribution by normalizing the coverage to sum to 1. The normalized observed coverage is termed the observed coverage distribution, which is comparable to the expected coverage distribution.

We use a slightly different definition of coverage from its classic meaning. We define the coverage of each transcript position to be the number of fragments starting at this position, while the classic definition considers the number of fragments spanning the position. We use the fragment start definition for calculating both the observed and the expected coverage distribution. The observed and the expected coverage are comparable if they are calculated using the same definition. Since the fragment length distribution is often assumed to be a Gaussian distribution with a smaller variance compared to the mean, the coverage distribution under the fragment start definition is approximately the same as the one under the classic definition plus a shift.

#### Definition 4.2

(Regional over-(under-)expression score). Given transcript *t* with length *l*, denote the expected coverage distribution as *exp*, and the observed coverage distribution as *obs*, the over-expression score of region [*a,b*] (1 ≤ *a* ≤ *b* ≤ *l*) is

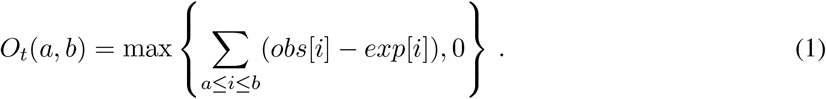

The under-expression score of region [*a, b*] is

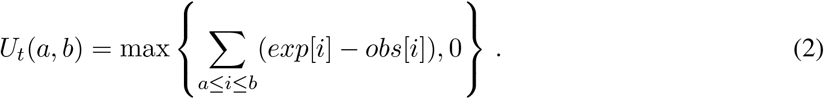

The over-expression and under-expression scores are defined as the probability difference between the observed coverage and the expected coverage distribution within region [*a, b*]. The probability difference represents the degree of inconsistency between the two distribution at the given region. The scores indicate the fraction of reads to take away (or add to) from the region in order for the two distributions to match each other.

#### Definition 4.3

(Transcript-level anomaly metric). For a transcript *t* with length *l*, the over-expression anomaly of the transcript is defined as

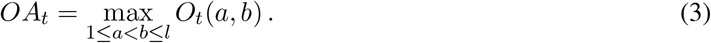

The under-expression anomaly of the transcript is defined as

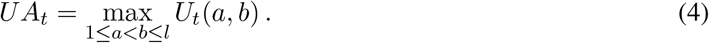

The transcript-level anomaly metric is defined by the largest over- or under-expression score across all continuous regions.

### 4.2 Probabilistic model for coverage distribution

The value of the anomaly metric cannot be directly used to indicate an anomaly because its value can be confounded by transcript abundances and the estimation error of the expected distribution. When there are only a few reads sequenced from the transcript, randomness in read sampling can dominate the observed distribution. Because of this, the observed distribution will have large fluctuations along the transcript positions, and thus appear to have large deviation from the expected distribution. In addition, when the estimation of the expected distribution is inaccurate, the difference between the two distribution can also be large. To address these two confounding factors, we model the relationship between the coverage distributions using a probabilistic framework and calculate the p-value of the anomaly metric. With the statistical significance of an anomaly score, we are able to distinguish between true quantification anomalies and randomness from known confounding factors.

We model the value of the anomaly metric probabilistically given the two confounding factors (Figure 7). We use the model to indicate the distribution of the anomaly metric under the null hypothesis that it is not a true anomaly. For the transcript abundance confounding factor, we assume the observed distribution is generated from the hidden expected distribution through a multinomial distribution parameterized by the given number of reads, *n*. For the estimation error of the expected distribution, we assume the error in the expected distribution is Gaussian. We model the true expected distribution with a hidden variable that is equal to the estimated distribution plus error. We further assume that the Gaussian estimation error is generally the same across all transcripts. In practice, transcripts have different lengths and the Gaussian error vectors differ relative to the lengths. We therefore bin the transcripts with similar lengths into the same number of bins, and estimate a shared mean shift parameter *μ* and covariance Σ for the transcripts with the same number of bins.

**Figure 7:**
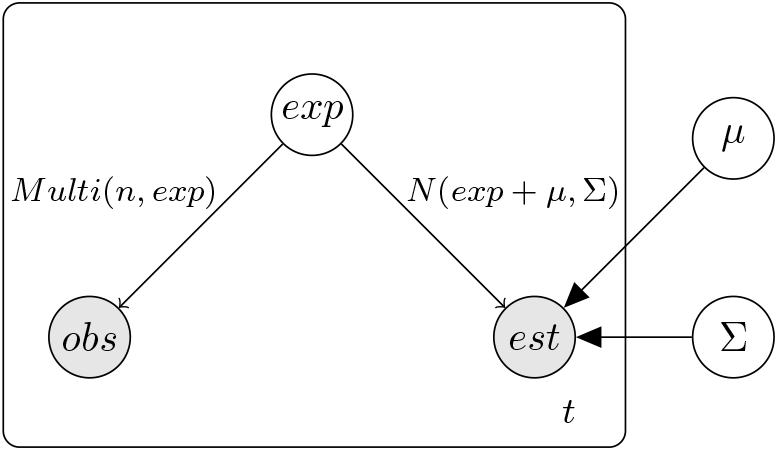
The probability relationship among the expected distribution, the observed distribution, and the estimator of the expected distribution. *exp* is the expected coverage, *obs* is the observed coverage, *est* is the estimation for the expected coverage. Here, *exp* is a hidden variable, while *obs* and est are observed. *obs* follows a multinomial distribution parameterized by the number of reads *n* and the expected coverage *exp. est* follows a Gaussian distribution with mean shift *μ* and covariance matrix Σ. We assume that the estimation errors of the expected coverage have the same pattern for all transcripts, and therefore *μ* and Σ are shared among all transcripts.

The variables and parameters of the model (Figure 7) can be retrieved or estimated as follows. *obs* refers to the observed distribution and can be retrieved from the quantification algorithm (Section 4.5). *est* refers to the estimation of the expected distribution, which is processed from the bias correction result of the quantification (Section 4.4). *exp* stands for the expected coverage distribution that is latent. *μ* and Σ in the probability could be estimated with a Bayesian estimator or maximum a priori (MAP) estimator with a likelihood function. Using subscript *t* to represent transcripts, the likelihood function is

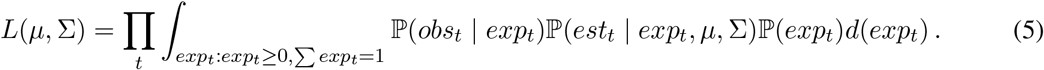

However, the above likelihood function does not have a closed form solution and may require using an expectation maximization (EM) approach for optimization, which is more than necessary. Instead, we estimate *μ* and Σ using the following approximation: the multinomial distribution for the observed coverage can be approximated by a Gaussian distributio*n* when the number of reads *n* is large enough:

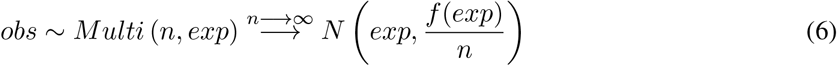

where *f* : ℝ^*m*^ → ℝ^*m×m*^ maps the *m*-dimension probability vector of the multinomial distribution into the covariance matrix of the approximating multi-variate Gaussian distribution. Therefore, the difference between *obs* and *est* can be approximated by the following Gaussian distribution

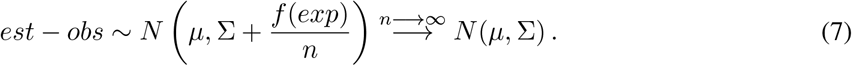

We therefore approximate *μ* and Σ by selecting transcripts with enough reads for each length group, and fit a Gaussian distribution to *est – obs* of the selected transcripts.

This probabilistic model serves as the null model that assumes the transcript is not an anomaly. That is, the model describes the distribution of the anomaly metric under the case where the deviation between the observed and the expected distribution is only due to the two confounding factors: read sampling randomness of sequencing and the estimation of expected distribution. When the deviation is so large that this null model cannot explain it, we attribute the deviation to an anomaly. To determine whether the deviation is so large that it is unlikely to be observed under the null model, the p-value is calculated, and the details of this calculation are explained in Section 4.3.

### 4.3 Statistical significance of the anomaly metric

The statistical significance of a value of the anomaly metric is the probability of observing an even larger anomaly value given the probabilistic model. Denote *O_t_*(*a, b*) and *U_t_*(*a, b*) to be the random variable of the regional over- and under-expression score of region [*a,b*], and denote *o_t_*(*a,b*) and *u_t_*(*a,b*) to be the corresponding observed values. Similarly, denote *OA_t_* and *UA_t_* to be the random variable of transcript-level anomaly score, and *oa_t_* and *ua_t_* to be the corresponding observed values. The p-values for a regional over- and under-expression score are

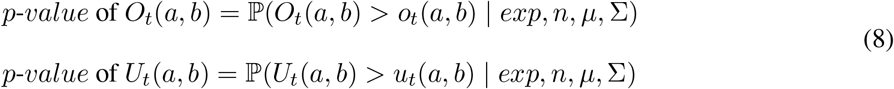

where *exp, n, μ* and Σ are defined as in Figure 7. The p-values for transcript-level over- and underexpression anomaly metric are

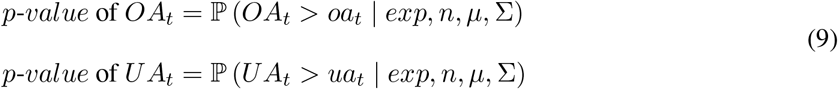

The statistical testing of transcript-level anomaly metric is more strict to the null hypothesis than the regional one, and tends to have a larger p-value. Given transcript *t* and the largest over-expression region [*i,j*], we have

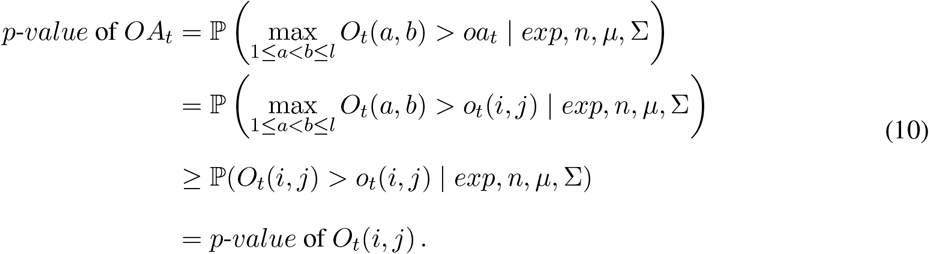

Conceptually, because the whole transcript contains multiple regions that may have a large over- (under-) expression score, it is easier to observe a large over- (under-) expression score when we look at all possible regions compared to when we focus on only one specific region. From the perspective of statistical testing, the p-value of *OA_t_* and *UA_t_* tend to be larger and less significant than those of *O_t_*(*a, b*) and *U_t_*(*a, b*) for any region [*a, b*]. Taking advantage of the different level of strictness about the null model, we use the significance of *O_t_* and *U_t_* for initial selection of anomalies to adjust read assignment (Section 4.6), and use the significance of *OA_t_* and *UA_t_* for final selection of anomalies within the unadjustable anomaly category.

The p-value of both anomaly metrics can be calculated empirically. Specifically, the hidden expected coverage can be sampled from the estimation using multi-variate Gaussian distribution, and the observed coverage can be sampled from the new hidden coverage using multinomial distribution. The null distribution for *O_t_*(*a, b*), *U_t_*(*a, b*), *OA_t_* and *UA_t_* can be generated using the sampled observed and hidden expected coverage. The empirical p-values is the portion of times that the anomaly scores exceed the observed valued in the null distribution.

We also derive a numerical approximation for the p-value of regional anomaly metric. Empirical p-value calculation requires sampling distributions from a multinomial or multi-variate Gaussian distribution multiple times, which takes a long time computationally. A numerical approximation without sampling can greatly reduce the calculation time. Denote the region as [*a, b*] and the current under-expression anomaly score as *v*. The significance of the over-(under-) expression score under regional null distribution is given by

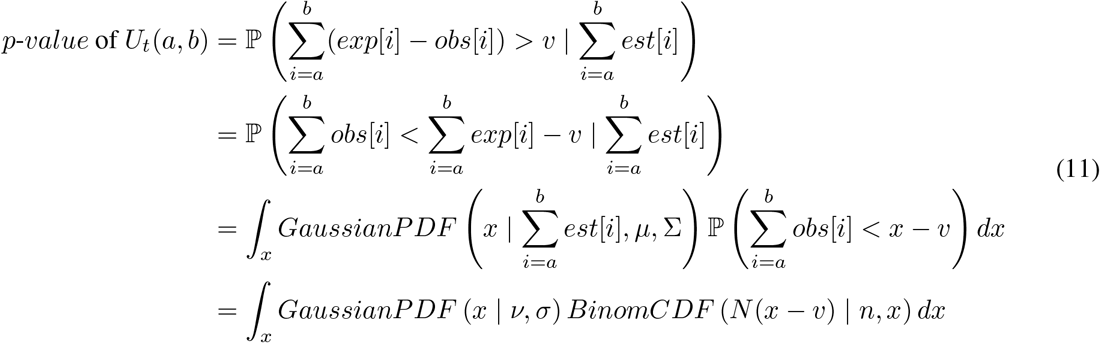

where 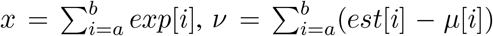, and 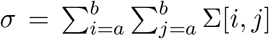. In numerical approximation, x takes value in a grid to sum the probabilities instead of computing the full integral. Since the regional anomaly metric focuses on a fixed region, the multinomial distribution can be collapsed into binomial distribution to represent the probability of generating a read from that region. The multi-variate Gaussian distribution can also be collapsed to a single-variate Gaussian distribution to present the expected estimation bias and variance of the region. With all multi-variate distributions collapsed into single-variate distributions, it is feasible to numerically calculate the integral in equation 11. In SAD, the p-value of the regional over- (under-) expression score is always calculated using the numerical approximation, while the p-value of the transcript-level anomaly is calculated empirically by sampling.

In practice, we do not calculate the p-value for transcripts with very low abundance. When the randomness of read sampling is very large, we simply assume that the p-value will be dominated by the randomness instead of incomplete reference transcriptome or quantification algorithm mistake. We only calculate a p-value for transcripts with average base pair coverage > 0.01. Using a threshold of 0.01 is equivalent to requiring that on average at least one read is sequenced for every 100 base pair.

Benjamini-Hochberg correction is used to control the rate of falsely discovered transcripts with regional or transcript-level expression anomaly. A threshold of 0.05 is used in regional anomaly score. For transcript-level anomalies, 0.01 is used as the threshold. The varied thresholds are set according to their separate purposes: regional anomalies are the initial candidates and do not need to be as precise; after read reassignment, the transcript-level anomalies are the final predictions of unadjustable anomaly and require precision.

### 4.4 Estimation of the expected distribution

The expected distribution is estimated for each transcript using the bias model from Patro et al. [7]. In the ideal case of sequencing, where the read is sampled randomly without any biases, the expected coverage is uniform along the positions of any transcript. However, in the real sequencing experiments, cDNA fragmentation and PCR amplification have preferences towards certain positional, sequence, and GC patterns, and the coverage is not expected to be uniform. The expected distribution is calculated to represent the probability of sampling a read at given position of given transcript. Salmon estimates the positional, sequence and GC biases by adjusting the uniform distribution based on the read mapping. There could be other biases affecting the expected distribution. However, other biases are not considered in the model, and thus the bias correction model is only an approximation for the expected distribution.

We processed the auxiliary output from Salmon to obtain the estimated expected distribution. The estimation for the expected distribution can also be calculated for other quantification software from bias correction model if corresponding output is available.

### 4.5 Observed distribution

The observed distribution is the actual read coverage for each transcript. It is calculated by counting the weighted number of reads at each position at given transcript after the weights are optimized by Salmon’s algorithm [7]. Specifically, when a read is multi-mapped to several transcripts, the weight represents the probability that the read is generated from the transcript.

### 4.6 Categorizing anomalies by re-assigning reads with linear programming

We categorize the causes of anomalies by re-assigning the reads and re-calculating the anomaly metric and its significance. Here, we only consider two causes: read assignment mistakes from the quantification algorithm and the incompleteness of the input reference.

We use linear programming (LP) to re-assign the reads. The LP formulation tries to use a linear combination of the expected distributions to explain the aligned reads. By explicitly using the expected coverage to re-distribute the observed number of reads, the deviation between the observed and the expected distribution after the re-distribution is naturally reduced. Accordingly, the anomaly score will decrease and the p-value will increase. We apply LP re-distribution separately for each gene since most reads are only multi-aligned across isoforms from the same gene.

To do this, we take the gene sequence to be the concatenation of all unique base pairs in its exons. The observed and the expected coverages are converted into gene-level coordinates. In this process, the observed coverage is not normalized and sums to the number of reads in the transcript assigned by Salmon [7]. Denote the set of transcripts of a gene by *T*, the expected distribution of transcript *t* ∈ *T* as *exp_t_*, and the observed read count vector is *obs_t_*. For transcript *t*_1_ and *t*_2_ within the same gene, *exp*_*t*1_, *exp*_*t*2_, *obs*_*t*1_ and *obs*_*t*2_ are of the same length. The LP for the re-assignment is

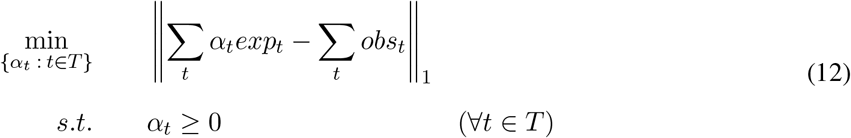

Variables *α_t_* stand for the expected number of expressed reads from transcript *t*. The actual number of reads re-assigned to transcript *i* at position *j* is 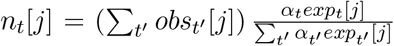. The actual total number of reads re-assigned to transcript *i* is Σ_*j*_ *n_t_* [*j*].

After adjusting read assignments by the LP, some of the regional anomalies become insignificant. Those are labeled “adjustable anomalies,” and the transcripts containing those regional anomalies are considered to have quantification algorithm error. To predict whether a transcript is affected by a novel isoform among the significant regional anomalies, we calculate the p-value of their transcript-level anomaly score and use Benjamini-Hochberg correction to control for the false positive labeling of anomalies for all transcripts. The transcripts with a significant transcript-level anomaly score are labeled as “unadjustable anomalies” and are predicted to be affected by novel isoform.

### 4.7 Minimizing the number of transcripts involved in read re-assignment

In practice, we try to keep the number of transcripts involved in the LP as small as possible. When the quantification of a transcript is good enough, re-assigning the reads may lead to a decrease of quantification accuracy. The correctness of the LP re-assignment largely depends on the accurate estimation of the expected distribution. However, the accuracy assumption of the expected distribution may not hold for all transcripts. An inaccurate estimation at some positions for one transcript can perturb the re-assignment result across all involved isoforms. The perturbation can be large when the coefficient matrix in the LP have a large condition number (called ill-conditioned), which tends to occur more often as the number of involved isoforms increases. The ill-condition will make the output very sensitive to a small change or error of the input distributions. To reduce the large perturbation problem in LP re-assignment, we only apply the LP reassignment on a small number of isoforms, and reset the other isoforms to the quantifier’s read assignment. The choice of isoforms is determined by the following principle: reducing the largest number of significant transcripts while at the same time minimizing the number of isoforms involved in the LP. An isoform is considered to be unnecessary in the LP if (1) it has an insignificant p-value and (2) after excluding it from LP the same set of significant isoforms remains insignificant under the re-assignment. The principle can be viewed as the removal of all unnecessary isoforms from read re-assignment.

To detect the unnecessary isoforms, we first identify the largest subset of significant transcripts that can become insignificant in re-assignment by initially running the LP using all transcripts. Then we exclude each insignificant transcript one by one from LP and test whether the exclusion retains the same subset of transcripts as insignificant. Labeling unnecessary isoforms requires iteratively running the LP optimization and the significance calculation. After all unnecessary isoforms are labeled, the iterative process is terminated.

## Acknowledgements

This research is funded in part by the Gordon and Betty Moore Foundation’s Data-Driven Discovery Initiative through Grant GBMF4554 to C.K., by the US National Science Foundation (CCF-1256087, CCF-1319998) and by the US National Institutes of Health (R01GM122935). This work was partially funded by The Shurl and Kay Curci Foundation. This project is funded, in part, under a grant (#4100070287) with the Pennsylvania Department of Health. The Department specifically disclaims responsibility for any analyses, interpretations or conclusions. The authors thank Dan DeBlasio and Rob Patro for helpful comments on this manuscript.

## Availability

The implementation of SAD is available at https://github.com/Kingsford-Group/sad.

## Authors’ contributions

CM and CK designed the method. CM implemented the algorithm and conducted the experiments. CM and CK drafted the manuscript. All authors read and approved the final manuscript.

