## Supplementary Text for "Detecting anomalies in RNA-seq quantification"

### Supplementary Material for
### Detecting RNA-seq quantification anomalies

C. Ma

C. Kingsford

February 5, 2019

#### 1 Simulation procedure

To mimic the real scenario where the target transcriptome contains novel transcripts outside reference transcriptome, we simulated target and reference transcriptome as follows. Using the gencode annotation [1], we randomly selected 200, 500, 1000, 1500 genes, remove one transcript per gene, and use the rest of the transcript sequences as reference transcripts. For the target transcriptome, we simulated 200, 500, 1000, 1500 fusion genes, added them to gencode transcript sequences, and use the combined full gencode transcripts and fusion sequences as the target transcriptome that generates RNA-seq data. In this case, the target transcriptome contains novel isoforms and fusion sequences compared to reference. We use both the protein-coding-only annotation and full annotation for removal and fusion simulation, to test both polyA RNA-seq and total RNA-seq techniques.

Reads are simulated using the target transcriptome by Polyester [2]. A count matrix is used as input in Polyester to denote the theoretical number of reads to be simulated for each transcript in the transcriptome. The count matrix is generated by quantifying RNA-seq datasets (GEUDAVIS, GM12878, K562) using Salmon [3] and the original gencode annotation.

With the simulation datasets, Salmon version 0.9.1 is used to quantify the reads against the simulated reference transcript.

#### 2 Running transcriptome assembly on simulated and real data

We ran Scallop version v0.9.8 on all simulated, GEUVADIS and Human Body Map samples, and set all parameters to their default. We also ran StringTie version 1.3.1c on all these samples, using the option "-G" for guiding the transcriptome assembly by the reference transcriptome. When guided by reference transcriptome, the precision of StringTie can be better than Scallop on some samples. We do not guide the Scallop assembly by reference transcriptome since it does not have the option. We use gffcompare [4] to compare the assembled transcripts with the reference transcript.

### 3 Supplementary figures

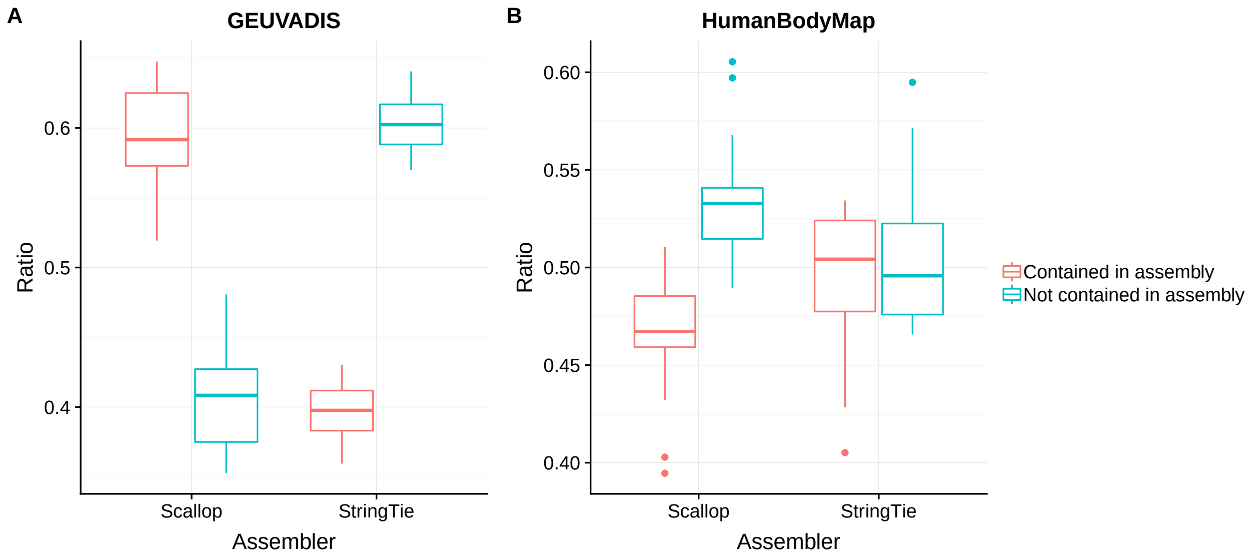


Figure S1: Proportion of the unadjustable-anomaly-containing genes that can (cannot) be detected by transcriptome assemblers. (A) For the GEUVADIS dataset, about 40% of the genes do not have corresponding novel isoforms predicted by Scallop, and about 60% of the genes do not have novel isoforms predicted by StringTie. (B) For the Human Body Map dataset, about 53% of novel-isoform-containing genes cannot be detected by Scallop, and about 50% of them cannot be detected by StringTie. The lower percentage of detection from StringTie in GEUVADIS dataset may be an effect of using the "Guided by reference" option.

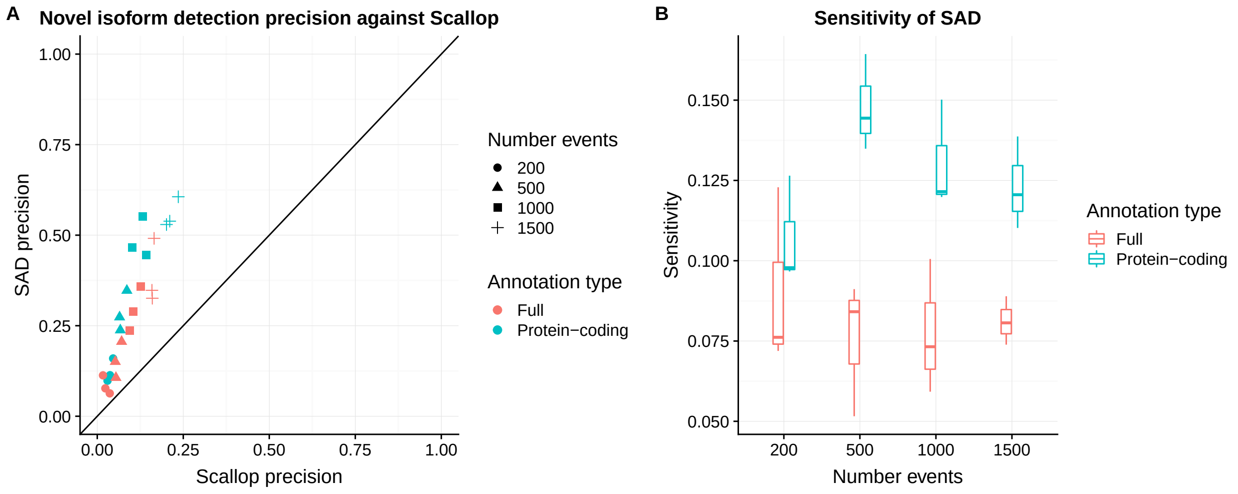


Figure S2: (A)Precision of novel isoform detection of SAD and Scallop. Point color and shape refers to different simulation settings. Precision is evaluated on the subset of novel isoforms that do not have novel splicing junctions, but only have alternative starting /ending sites, or a different combination of existing splicing junctions in the reference annotation. (B)Sensitivity of the unadjustable anomalies of SAD. Most of the simulated novel isoforms do not affect the coverage significantly enough to be detected by SAD. Annotation type indicates whether the Salmon quantification uses the full set of known transcripts or the protein-coding-only transcripts.

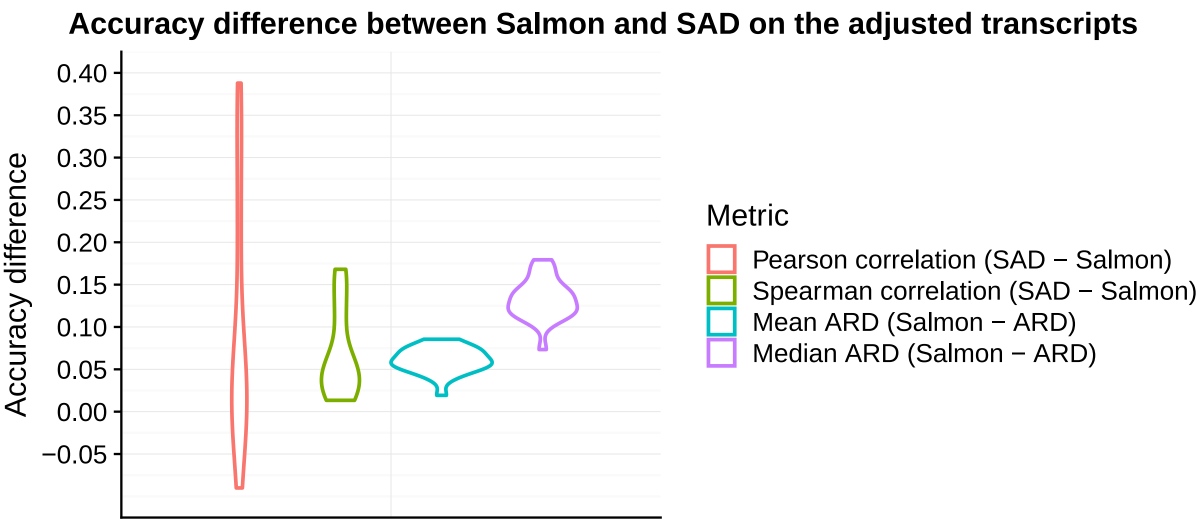


Figure S3: Comparing the accuracy of SAD adjusted quantification with the accuracy of Salmon's original result on simulation data. Accuracy is evaluated on the subset of transcripts involved in read re-assignment of SAD. Two similarity measures, Pearson correlation and Spearman correlation, and two dissimilarity measures, mean ARD and median ARD, are used as the measure of accuracy. For all four measurements, values above 0 indicates that SAD is more accurate than Salmon. Under the two similarity measure, the SAD adjusted quantification is more accurate than Salmon for some of the simulated datasets, and achieves similar accuracy for the rest. Under the two dissimilarity measure, SAD is more accurate than Salmon for all datasets.
